# Macrophage-Dependent Trafficking and Remodeling of the Basement Membrane-Interstitial Matrix Interface

**DOI:** 10.1101/364422

**Authors:** Julian C. Bahr, Stephen J. Weiss

**Affiliations:** Cancer Biology Graduate Program, University of Michigan, Ann Arbor, MI 48109; Division of Molecular Medicine and Genetics, University of Michigan, Ann Arbor, MI 48109; Department of Internal Medicine, University of Michigan, Ann Arbor, MI 48109; Life Sciences Institute, University of Michigan, Ann Arbor, MI 48109; Rogel Cancer Center, University of Michigan, Ann Arbor, MI 48109

## Abstract

Macrophages infiltrate and remodel extracellular matrix barriers during development and disease. Bahr and Weiss demonstrate that human macrophages remodel native basement membrane barriers in an MT1-MMP-dependent fashion while retaining the ability to traverse tissue barriers by actomyosin-dependent forces alone.

**Abstract:** Macrophages dominate inflammatory environments where they modify the extracellular matrix by mobilizing complex repertoires of proteolytic enzymes. Nevertheless, the dominant proteinases used by macrophage as they confront physiologic tissue barriers remain undefined. Herein, we have characterized the molecular mechanisms that define human macrophage-extracellular matrix interactions *ex vivo*. Resting and immune-polarized macrophages are shown to proteolytically remodel basement membranes while infiltrating the underlying interstitial matrix. In an unbiased screen to identify key proteases, we find that the macrophage metalloproteinase, MT1-MMP, is the dominant effector of basement membrane degradation and invasion. Unexpectedly, macrophages can alternatively use actomyosin-dependent forces to transmigrate native basement membrane pores that provide cells with proteinase-independent access to the interstitial matrix. These studies not only identify MT1-MMP as a key proteolytic effector of extracellular matrix remodeling by human macrophages, but also define the invasive strategies used by macrophages to traverse physiologic tissue barriers.

## Introduction

Macrophages infiltrate and remodel the extracellular matrix (ECM) of native tissues under a wide variety of physiologic and pathologic conditions, ranging from post-parturition mammary gland involution to metastasis (Lewis et al., 2016; Wynn, Chawla, & Pollard, 2013; Wynn & Vannella, 2016). In mediating these diverse effects, macrophages assume an array of differentially activated or polarized states that allow them to either degrade or repair the ECM (Afik et al., 2016; P. J. Murray et al., 2014; Talmi-Frank et al., 2016). Regardless of their activation state, however, macrophages interact with at least one of two distinct ECM compartments, i.e., the basement membrane or the interstitial matrix (Rowe & Weiss, 2008, 2009). As a specialized form of ECM, the basement membrane subtends all epithelial and endothelial cell layers, but also surrounds adipocytes, pericytes, nerves, and vascular smooth muscle cells (Fidler et al., 2017; Rowe & Weiss, 2008). Despite ranging in thickness from only 100–300 nm, basement membranes are mechanically rigid barriers in almost all tissues, largely owing to a covalently cross-linked network of tightly intertwined type IV collagen fibers that non-covalently associate with a laminin meshwork as well as a complex mix of more than 70 other components (Halfter et al., 2015; Randles, Humphries, & Lennon, 2017). In turn, the underlying interstitial matrix is dominated by an interwoven composite of fibrillar type I/III collagen, elastin, glycoproteins, proteoglycans, and glycosaminoglycans (Rowe & Weiss, 2009).

In confronting the basement membrane-interstitial matrix continuum *in vivo*, current evidence suggests normal as well as neoplastic cells remodel the ECM interface in order to drive tissue-invasive activity (Chang, Thakar, & Weaver, 2017; Kelley, Lohmer, Hagedorn, & Sherwood, 2014; Rowe & Weiss, 2008; Sabeh, Shimizu-Hirota, & Weiss, 2009).

To date, however, efforts to characterize human macrophage-ECM interactions have largely been confined to the use of artificial matrix constructs that lack the critical structural organization and mechanical properties that characterize the ECM *in vivo* (Cougoule et al., 2012; Halfter et al., 2015; Jevnikar et al., 2012; Daniel Hargbøl Madsen et al., 2017; Randles et al., 2017; Rowe & Weiss, 2008, 2009; Starnes et al., 2014; Van Goethem, Poincloux, Gauffre, Maridonneau-Parini, & Le Cabec, 2010; Wiesner, Le-Cabec, El Azzouzi, Maridonneau-Parini, & Linder, 2014). Hence, despite the fact that macrophages can mobilize a complex repertoire of proteolytic enzymes belonging to the aspartyl-, serine-, cysteine- and metallo-proteinase families, which, if any, of these systems participate in tissue remodeling and invasion remains undefined (Akkari et al., 2014; Nathan & Ding, 2010; Newby, 2016; Sevenich & Joyce, 2014; Vérollet et al., 2011). To this end, we now characterize the molecular mechanisms that underlie macrophage-dependent remodeling of native ECM barriers. Unlike other cell populations characterized to date, we find that human macrophages display a unique hybrid ability to penetrate native tissues by either mobilizing the membrane-anchored matrix metalloproteinase, MT1-MMP, that serves to dissolve intervening matrix barriers or alternatively, generating actomyosin-dependent mechanical forces that drive a shape shifting phenotype that permits invasion to proceed independently of matrix-degradative activity.

## Results

### Primary human macrophages remodel native basement membrane

Following gentle decellularization of native mesenteric sheets, 3-dimensional (3D) reconstructions of immunofluorescent and second harmonic generation images allow visualization of a reflected basement membrane bilayer that ensheaths an intervening interstitial matrix (Figure 1A,B and Video 1–2) (Hotary, Li, Allen, Stevens, & Weiss, 2006; Witz et al., 2001). At higher magnification, *en face* confocal images of excised tissue incubated with anti-laminin or anti-type IV collagen antibodies highlight the confluent nature of the basement membrane sheets (Figure 1C). By imaging through the labeled tissue, orthogonal xz and yz reconstructions permit visualization of the apical and basal basement membrane layers that are separated by the ~50 μm-thick (unstained) interstitial matrix (Figure 1C). Given that the mechanical integrity of basement membranes is largely defined by a variable number of intermolecular sulfilimine bonds formed between the C-terminal domains of opposing type IV collagen trimers (Figure 1D) (Fidler, Boudko, Rokas, & Hudson, 2018), we sought to determine the relative frequency of these covalent cross-links in the isolated basement membranes. Following digestion with bacterial collagenase, the C-terminal domains of type IV collagen molecules (termed NC1 domains) remain associated as either non-covalently or covalently-associated dimers (Boudko, Danylevych, Hudson, & Pedchenko, 2018; Fidler et al., 2014). However, following SDS-PAGE, only the covalently cross-linked dimers remain intact, whereas non-covalently cross-linked dimers dissociate into monomers (Figure 1E) (Boudko et al., 2018). In the specific case of the peritoneal basement membrane, analyses of NC1 dimer structure demonstrate that its type IV collagen network is dominated by covalent cross-links (Figure 1E), thereby highlighting the fact that tissue-invasive cell populations confront a mechanically stable barrier.

**Figure 1.**
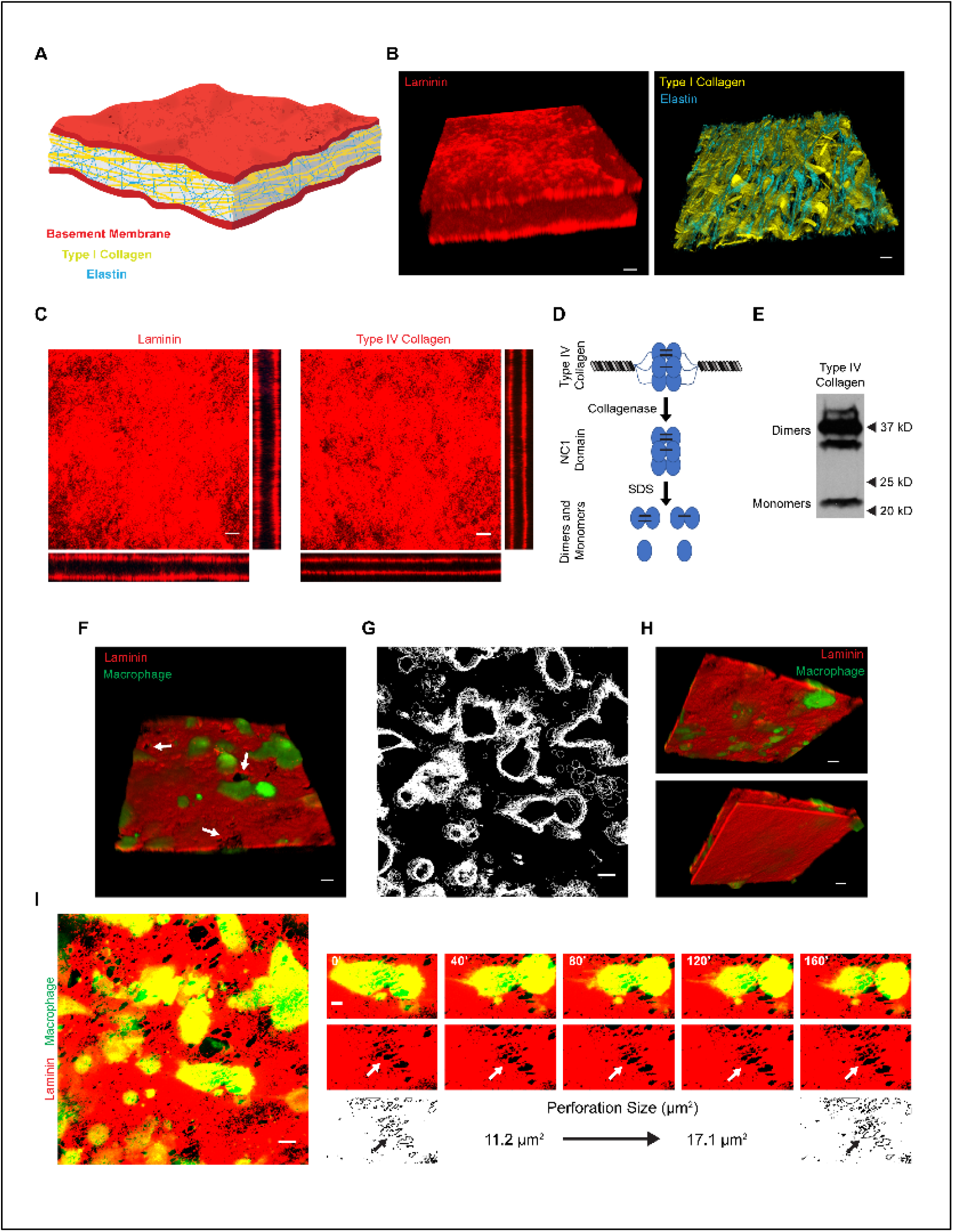
Human macrophage interactions with native basement membrane. **(A)** Schematic illustration of the mesentery extracellular matrix. **(B)** 3D confocal reconstructions of laminin (red, left panel) and elastin (blue, right panel), with second harmonic generation of type I collagen (yellow, right panel) in rat mesentery constructs. **(C)** *En face* and orthogonal immunofluorescence of laminin and type IV collagen. **(D)** Schematic of type IV collagen dimer-monomer content analysis. After collagenase digestion of type IV collagen, the hexameric NC1 domain remains intact. The hexamer can be dissociated via non-reducing SDS-PAGE into sulfilimine-crosslinked dimers and non-crosslinked monomers. **(E)** Type IV collagen dimer-monomer content analysis as determined by Western blotting. **(F)** 3D confocal reconstruction of human macrophages (green) atop the apical face of a basement membrane (red) with adjacent perforations (arrow) after 48 h. **(G)** Overlay of macrophage outlines captured every 10 minutes for 160 minutes. **(H)** 3D reconstruction from (F) rotated 180° showing the apical basement membrane surface (top panel) and the reflected basal face (bottom panel). **(I)** Immunofluorescence of the apical basement membrane layer and macrophages (left panel) with a macrophage actively expanding a perforation in the basement membrane (small panels, arrows) from an area of 11.2 μm^2^ to 17.1 μm^2^ over 160 minutes. Bars: 20 μm (B, F, H); 10 μm (C, G, I left panel); 5 μm (I right panel).

Given the compositional and structural integrity of our *ex vivo* tissue construct, we next sought to characterize the nature of its interactions with primary human macrophages. As such, carboxyfluorescein diacetate succinimidyl ester (CFSE)-labeled monocyte-derived macrophages were cultured atop basement membranes that were pre-labeled with fluorescently-tagged anti-laminin antibodies in the presence of F_c_ receptor blocking reagents to prevent direct interactions between the macrophages and the antibody-coated surface (Güç, Fankhauser, Lund, Swartz, & Kilarski, 2014; Kilarski et al., 2013). After a 48 h time period, the macrophage-tissue constructs were imaged for 160 mins using real-time spinning disc confocal microscopy. As shown, macrophages (green) are found adherent to the basement membrane in association with the appearance of distinct 5–10 μm diameter perforations in the labeled matrix (Figure 1F, arrows). While real-time imaging of macrophage membrane contours over this timespan demonstrates only small changes in lateral cell spreading (Figure 1G), cells are observed that are actively traversing the apical face of the *ex vivo* construct with macrophage membrane protrusions found breaching the basement membrane surface (Figure 1H and Video 3). Indeed, under high magnification, real-time imaging of a single basement membrane pore in association with an overlying macrophage over this time span demonstrates an increase in perforation size from ~11 μm^2^ to ~17 μm^2^ in the absence of noticeable changes in the fluorescent intensity of the pore edge (Figure 1I and Video 4), a finding consistent with the active proteolytic remodeling of the cell-matrix interface. Hence, by 48 h, human macrophages are able to remodel native basement membranes while breaching the surface with invasive membrane protrusions.

### Inflammatory stimuli alter the basement membrane remodeling potential of human macrophages

Given that macrophages serve discrete functions during the initiation and resolution of inflammatory responses (Wynn & Vannella, 2016), we sought to characterize the effect of immune polarizing stimuli on basement membrane remodeling. Consistent with recent studies, macrophages stimulated with purified E. coli lipopolysaccharide (LPS) upregulate *TNFα* and downregulate *MRC1* transcript levels (Figure 2A) (Martinez et al., 2013; P. J. Murray et al., 2014). Conversely, polarizing macrophages with the cytokine, IL-4, downregulates *TNFα* and upregulates *MRC1* transcript levels (Figure 2A) (Martinez et al., 2013; P. J. Murray et al., 2014). As such, variably polarized macrophages were cultured atop *ex vivo* tissue constructs before staining with anti-laminin antibodies to assess basement membrane integrity. After a 6-day culture period, unstimulated macrophages are observed atop the apical basement membrane in association with large numbers of ~10 μm diameter perforations (Figure 2B, arrowheads). Over this timeframe, orthogonal reconstructions demonstrate that macrophages not only remodel the basement membrane interface, but also proceed to infiltrate the underlying interstitial matrix (Figure 2B, xz and yz images). Of note, a subset of the tissue-invasive macrophages also begin to traverse the opposing reflected basement membrane, demonstrating that the remodeling program occurs regardless of basement membrane symmetry (Halfter, Candiello, et al., 2013) (Figure 2B arrowheads).

**Figure 2.**
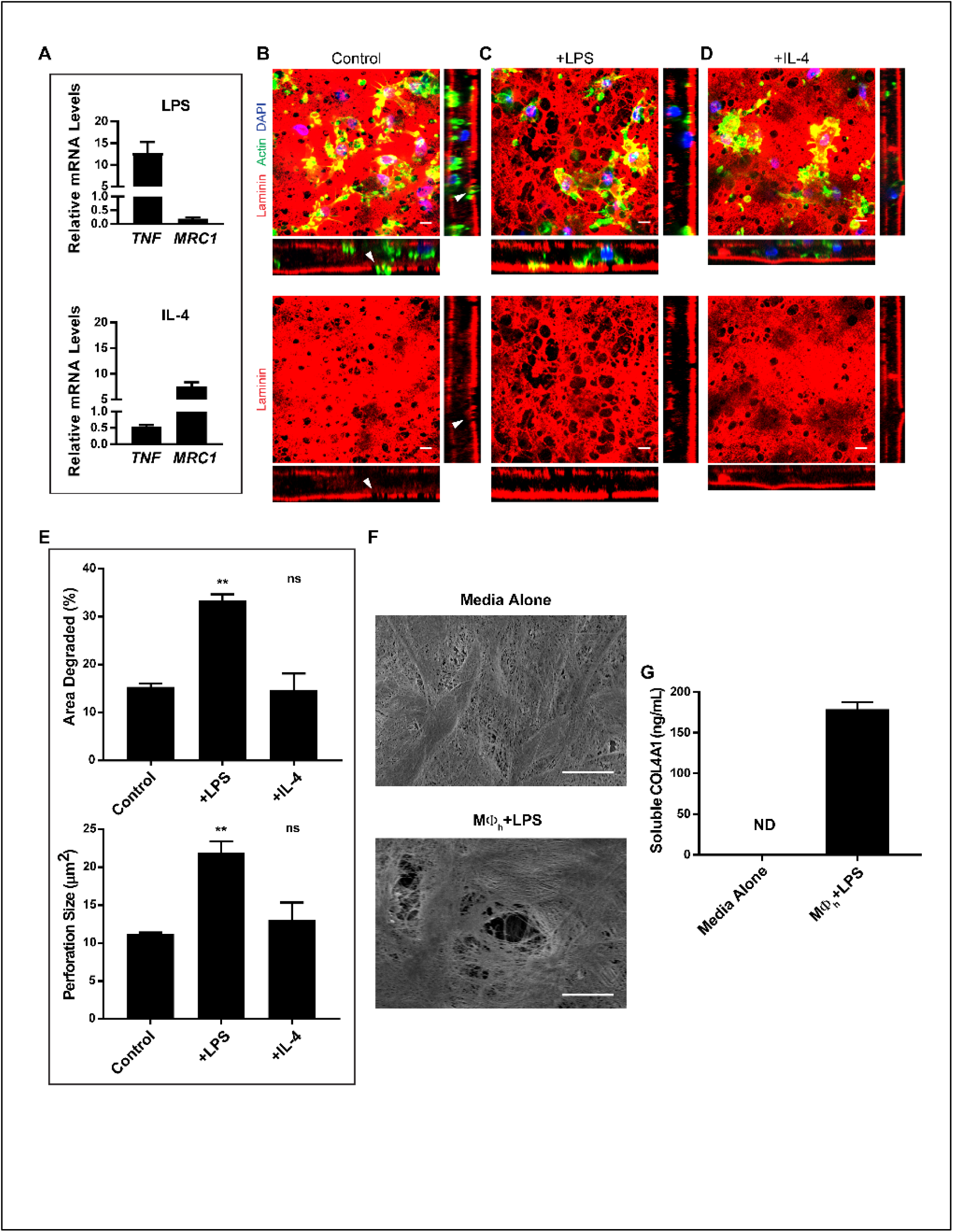
Polarized human macrophage-dependent remodeling of the basement membrane. **(A)** Transcript expression of immune response genes was analyzed by qPCR in human macrophages polarized with LPS (1 μg/mL) or recombinant human IL-4 (20 ng/mL). Results are expressed as mean ± SEM, n = 3, fold-change relative to control. **(B)** Immunofluorescence of unstimulated macrophages cultured for 6 days on the basement membrane surface. Orthogonal reconstructions reveal cells that infiltrated the interstitial matrix and perforated the basal basement membrane (arrows). **(C)** Immunofluorescence of macrophages polarized with LPS (1 μg/mL) or **(D)** recombinant human IL-4 (20 ng/mL). Images shown in (B-D) are representative of three replicates. **(E)** Quantification of the area of basement membrane degraded and basement membrane perforation size as analyzed by ImageJ pixel analysis of each condition from (B-D). Results are expressed as mean ± SEM; (**) P < .001; ns, not significant; n = 3. **(F)** Scanning electron micrograph of basement membrane stripped of cells either after culture with medium alone or with LPS-polarized macrophages for 6 days. **(G)** Quantification of soluble type IV collagen detected in cell-free media on day 3 of (F); results are expressed as mean ± SEM, n = 4. Bars: (B-D, F) 10 μm.

In a manner similar to unstimulated macrophages, LPS-polarized cells likewise remodel the basement membrane, but the percent surface area degraded increases ~2-fold as does the average size of the perforations (Figure 2C,E). Interestingly, while IL-4-dependent polarization is commonly linked to a tissue remodeling phenotype (Daniel H. Madsen et al., 2013; Mantovani, Biswas, Galdiero, Sica, & Locati, 2013), these macrophages fail to affect remodeling beyond that observed with unstimulated macrophages (Figure 2D,E).

Given that LPS-stimulated macrophages mount the most robust remodeling program, we used these cells to characterize the dynamic characteristics underlying the formation of basement membrane perforations. Macrophages actively engaged in the remodeling of adjacent perforations adopt two distinct morphologies, i.e., either encircling the border of the perforation without extending obvious protrusions into the cavity or sending small cell processes through the perforation into the underlying interstitial matrix (Video 5 and 6). Coincident with this activity, macrophages can be observed to ‘tug’ on the underlying basement membrane with force sufficient to contort the matrix (Video 6). As basement membrane perforations can be generated – or enlarged – as a function of reversible mechanical distortions (Kelley et al., 2014), we assessed basement membrane structure by SEM following the 6-day culture period. As shown, clearly demarcated perforations that are ~10 μm diameter can readily be found in macrophage-exposed, but not control, constructs (Figure 2F). In tandem with basement membrane denudation, type IV collagen is solubilized as assessed by ELISA (Figure 2G).

While recent studies have highlighted the ability of human and mouse macrophages to respond to specific inflammatory stimuli in transcriptionally and phenotypically distinct fashion (Martinez et al., 2013; Seok et al., 2013), we find that mouse bone marrow-derived macrophages (BMDM) also perforate the basement membrane in response to LPS and IL-4 polarization, as well as infiltrate the interstitial compartment (Figure 3A). Under these conditions, LPS-stimulated BMDM remodel an area three times larger than unstimulated or IL-4 stimulated BMDM, and form modestly, although not significantly, larger perforations (Figure 3C). Interestingly, mouse multinucleated giant cells are occasionally formed in response to IL-4 (McNally & Anderson, 1995), but they express only minimal basement membrane remodeling activity and fail to invade the interstitial matrix (Figure 3B). Taken together, these data demonstrate that human as well as mouse macrophages proteolytically remodel and actively transmigrate native tissue barriers through processes responsive to immune polarization.

**Figure 3.**
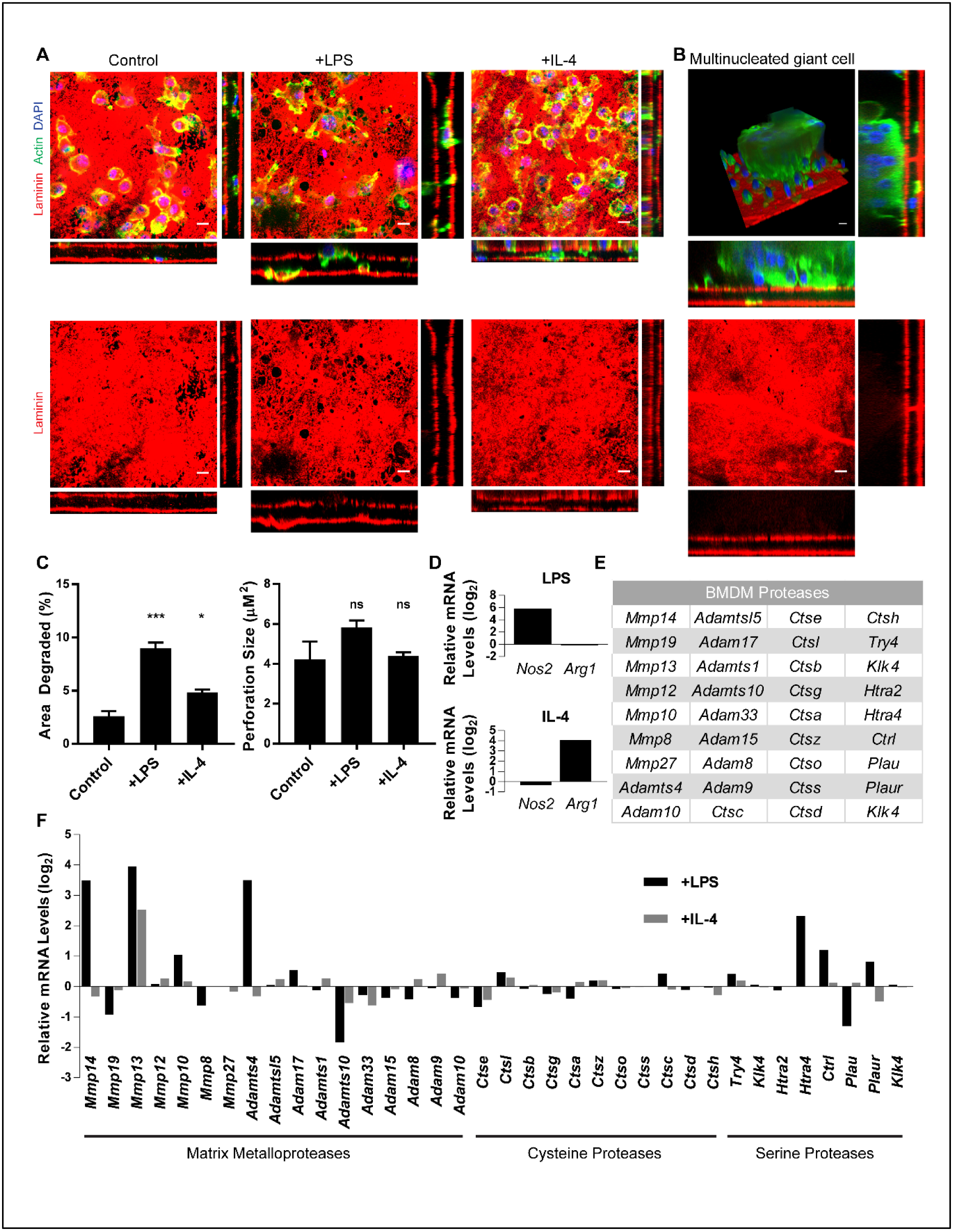
Polarized mouse macrophages express a suite of proteases. **(A)** Immunofluorescence of mouse BMDM cultured for 6 days on basement membrane constructs in the presence of media alone, LPS (1 μg/mL), or recombinant mouse IL-4 (20 ng/mL). Images shown are representative of three replicates. **(B)** 3D, *en face*, and orthogonal images of a multinucleated giant cell formed in response to IL-4. **(C)** Quantification of the area of basement membrane degraded and basement membrane perforation size as analyzed by ImageJ pixel analysis of each condition from (A). Results are expressed as mean ± SEM; (***) P < .0001, (*) P < .01; ns, not significant; n = 3. **(D)** Microarray data for two biological replicates of mouse BMDM left unstimulated, polarized with LPS (1 μg/mL), or polarized with recombinant mouse IL-4 (20 ng/mL) for 24 h. Relative expression levels of mouse-specific immune response genes (D), proteases with an absolute gene expression value of at least 2^4^ **(E)**, and the relative expression of those proteases in response to LPS and IL-4 **(F)** are presented. (D, F) are on a log_2_ scale.

### Macrophage polarization and BM remodeling correlates with protease expression

As both mouse and human macrophages display similar matrix-remodeling phenotypes, we sought to first use mouse BMDM as a genetically-modifiable system to identify the underlying mechanisms responsible for basement membrane remodeling. To this end, we transcriptionally profiled mouse BMDM after a 24 h culture period under either unstimulated, LPS-stimulated or IL-4-stimulated conditions. As expected, the upregulation of mouse-specific polarization markers, *Nos2* and *Arg1* (Gundra et al., 2014; P. J. Murray et al., 2014), correlated with LPS and IL-4 stimulation, respectively (Figure 3D). In addition, a large number of proteases belonging to the metalloproteinase, cathepsin and serine proteinase family thought to be essential for ECM remodeling are expressed under these conditions (Figure 3E) (Fleetwood et al., 2014; Jevnikar et al., 2012; Daniel H. Madsen et al., 2013; M. Y. Murray et al., 2013). Of note, however, only a relatively small number of these proteases are differentially expressed in response to LPS or IL-4, with a smaller subset of these enzymes altering their transcript levels in a pattern that correlated with the matrix-remodeling phenotype, including the metalloproteases, *Mt1-mmp* and *Adamts4;* serine proteases, *Htra4* and *Ctrl;* and serine protease receptor, *Pluar* (Figure 3F).

### Matrix metalloproteases are required for basement membrane remodeling

Cognizant of the fact that correlative changes in transcripts level may – or may not – correlate with matrix degradation activity, we next sought to identify effector proteases responsible for matrix remodeling by culturing BMDM atop tissue explants in the presence of broad spectrum inhibitors directed against cysteine, serine, or metalloprotease family members (Fleetwood et al., 2014; Hotary et al., 2006; Pflicke & Sixt, 2009; Van Goethem et al., 2010; Wolf et al., 2013). Despite the expression of multiple proteases by LPS-stimulated BMDMs, the addition of high concentrations of validated cysteine or serine protease inhibitors fail to inhibit basement membrane remodeling to a significant degree (Figure 4A) (Punturieri et al., 2000; Reddy, Zhang, & Weiss, 1995). In contrast, the metalloprotease inhibitor, BB-94, that targets MMP, ADAM and ADAM-TS family members (Baker, Edwards, & Murphy, 2002; Seals & Courtneidge, 2003) significantly blocks basement membrane degradation without affecting macrophage-basement membrane adhesion or cell viability (Figure 4A and data not shown). To further narrow the number of candidate proteases, we took advantage of the fact that endogenous protease inhibitors, known as tissue inhibitor of metalloproteinases (TIMPs), can be used to preferentially block the proteolytic activity of secreted versus membrane-anchored MMPs (Brew & Nagase, 2010; English et al., 2006; Hotary et al., 2006). In the presence of TIMP-1, a more specific inhibitor of secreted MMPs (English et al., 2006; Sabeh, Li, Saunders, Rowe, & Weiss, 2009), the remodeling program is unaffected (Figure 4A). By contrast, TIMP-2, an endogenous inhibitor of both secreted and type I membrane-anchored MMPs (English et al., 2006; Hotary et al., 2006; Sabeh, Li, et al., 2009), abrogates basement membrane degradation completely (Figure 4A). As BB-94 and TIMP-2 are the only inhibitors that effectively block basement membrane degradation, these results support the conclusion that a membrane-type MMP (MT-MMP) is likely the sole protease required for basement membrane remodeling (Figure 4B).

**Figure 4.**
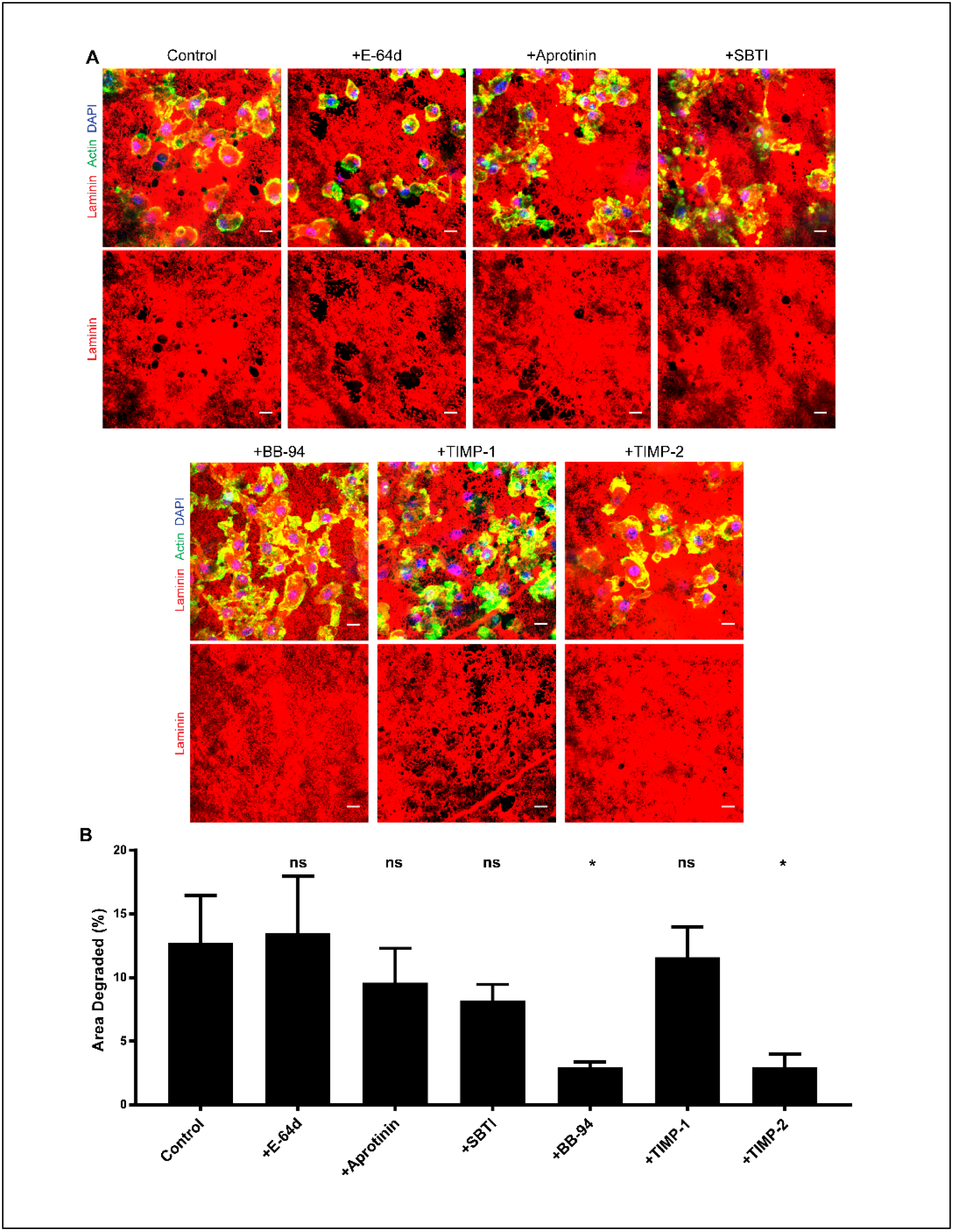
Mouse macrophages require matrix metalloproteinases for basement membrane remodeling. **(A)** Macrophages were cultured atop basement membrane constructs for 6 days with LPS (1 μg/mL) in the absence or presence of inhibitors directed against cysteine proteinases (100 μM E-64d), serine proteinases (100 μg/mL aprotinin; 100 μg/mL soybean trypsin inhibitor, SBTI), matrix metalloproteinases (5 μM BB-94), 12.5 μg/mL TIMP-1, or 5 μg/mL TIMP-2. Images are representative of three replicates. Bars: 10 μm. **(B)** Quantification of the area of basement membrane degraded as analyzed by ImageJ pixel analysis under each set of conditions. Results are expressed as mean ± SEM; (*) P < .01; ns, not significant; n = 3.

### MT1-MMP is the dominant effector responsible for macrophage-mediated remodeling of the basement membrane

While at least four members of the MT-MMP family are sensitive to TIMP-2 (i.e. MT1-MMP, MT2-MMP, MT3-MMP and MT5-MMP)(English et al., 2006; Rowe & Weiss, 2009), transcriptional profiling of LPS-stimulated macrophages identified MT1-MMP as the sole membrane-anchored MMP expressed under these conditions (Figure 3E,F). Given that the increase in MT1-MMP transcript levels most closely correlated with the basement membrane remodeling phenotype, we confirmed by Western blot and qPCR that MT1-MMP is significantly upregulated following polarization with LPS (Figure 5A). As such, to directly define the impact of MT1-MMP on the matrix remodeling program, BMDM were prepared from *Mt1-mmp^−/−^* mice and cultured atop native explants. Underlining an essential requirement for MT1-MMP in basement membrane remodeling, *Mt1-mmp^-/-^* BMDM fail to display matrix-degradative activity under basal, LPS-, or IL-4-stimulated conditions (Figure 5B,D), despite maintaining identical expression of non-targeted cysteine, serine, and metallo-proteases (data not shown). Importantly, following transduction of *Mt1-mmp^−/−^* macrophages with an MT1-MMP/mCherry-tagged construct (Sakurai-Yageta et al., 2008), basement membrane perforations materialize coincident with macrophages extending MT1-MMP/mCherry-positive protrusions into the underlying interstitial stroma (Figure 5C,D).

**Figure 5.**
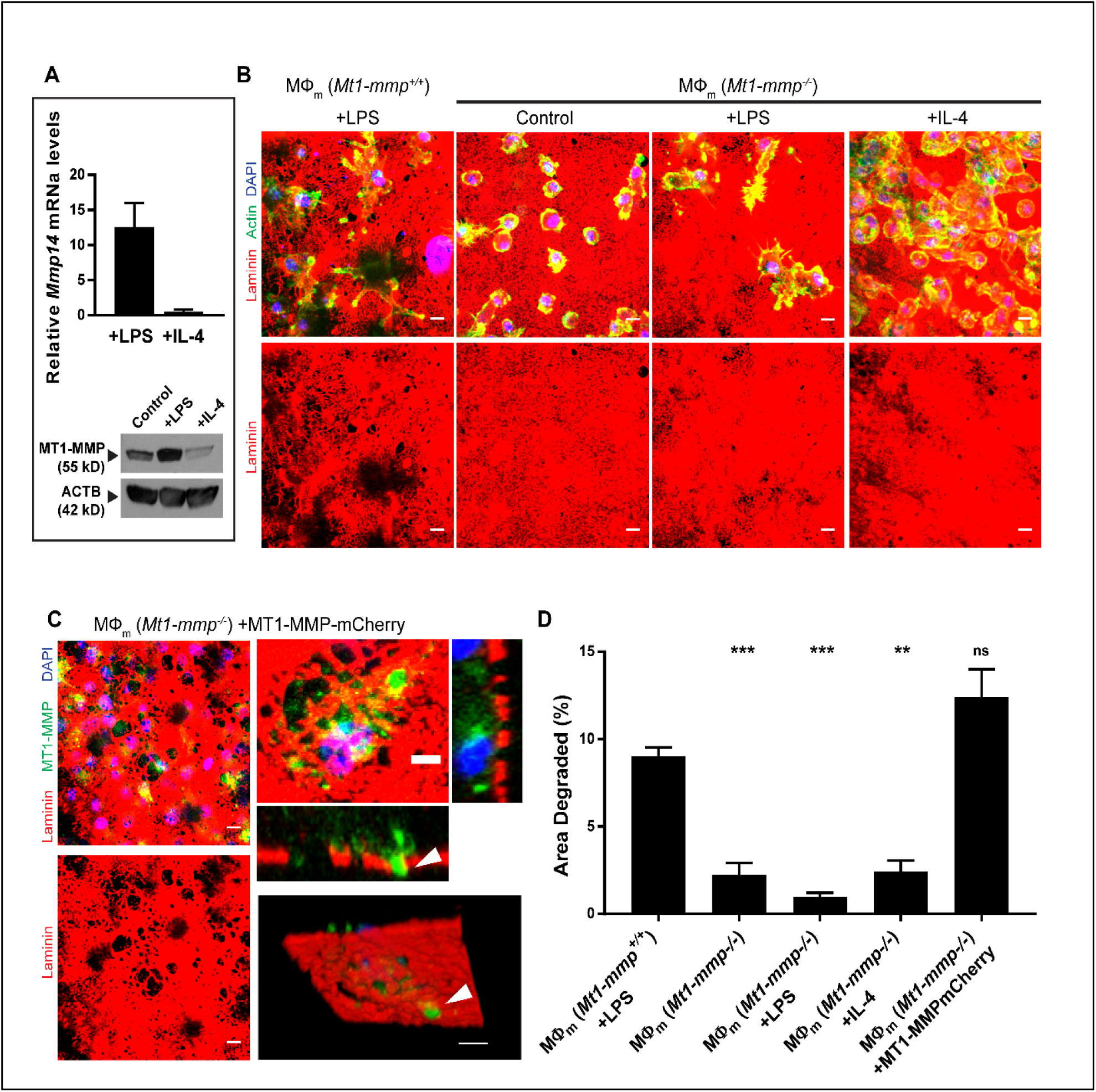
MT1-MMP-dependent mouse BMDM-mediated basement membrane remodeling. **(A)** Relative MT1-MMP expression in mouse BMDM left unstimulated or polarized with LPS (1 μg/mL), or recombinant mouse IL-4 (20 ng/mL), as determined by qPCR (top panel) or Western blot (bottom panel). **(B)** Immunofluorescence basement membranes exposed to either LPS-polarized MT1-MMP^+/+^ mouse BMDM or unstimulated, LPS-, and IL-4-polarized MT1-MMP^−/−^ mouse BMDM. **(C)** Immunofluorescence of MT1-MMP^−/−^mouse BMDM transduced with a lentiviral MT1-MMP-mCherry vector (pseudo-colored green) for 48 h before culture on the basement membrane construct (pseudo-colored red). MT1-MMP-mCherry-positive protrusions are localized to basement membrane perforations (arrowheads). Images shown in (B-C) are representative of three replicates. Bars: (B; C, left panels) 10 μm, (C, right panels) 5 μm. **(D)** Quantification of the area of basement membrane degraded as analyzed by ImageJ pixel analysis under each set of conditions from (B-C). Results are expressed as mean ± SEM; (***) P < .0001, (**) P < .001; ns, not significant; n = 3.

To next determine whether the mouse MT1-MMP-dependent regulation of basement membrane remodeling can be extended to human macrophages, we first confirmed that primary human macrophages express MT1-MMP and upregulate its expression following LPS polarization (Figure 6A). However, while LPS increased *MT1-MMP* transcript levels, protein expression remains largely unchanged (Figure 6A). Nevertheless, as assessed by confocal imaging, while endogenous MT1-MMP was found to localize in permeabilized cells to the peri-nuclear ER/Golgi region as well as trafficking vesicles throughout the cell, under non-permeabilized conditions, the levels of cell surface-associated MT1-MMP increase in response to LPS polarization (Figure 6B). Consistent with these findings, when human macrophages are cultured atop the basement membrane in the presence of BB-94 or a monoclonal antibody directed against the catalytic domain of MT1-MMP (Ager et al., 2015; Devy et al., 2009), matrix degradation is almost completely ablated (Figure 6C-F). Hence, MT1-MMP is required for both mouse and human macrophage-mediated basement membrane degradation.

**Figure 6.**
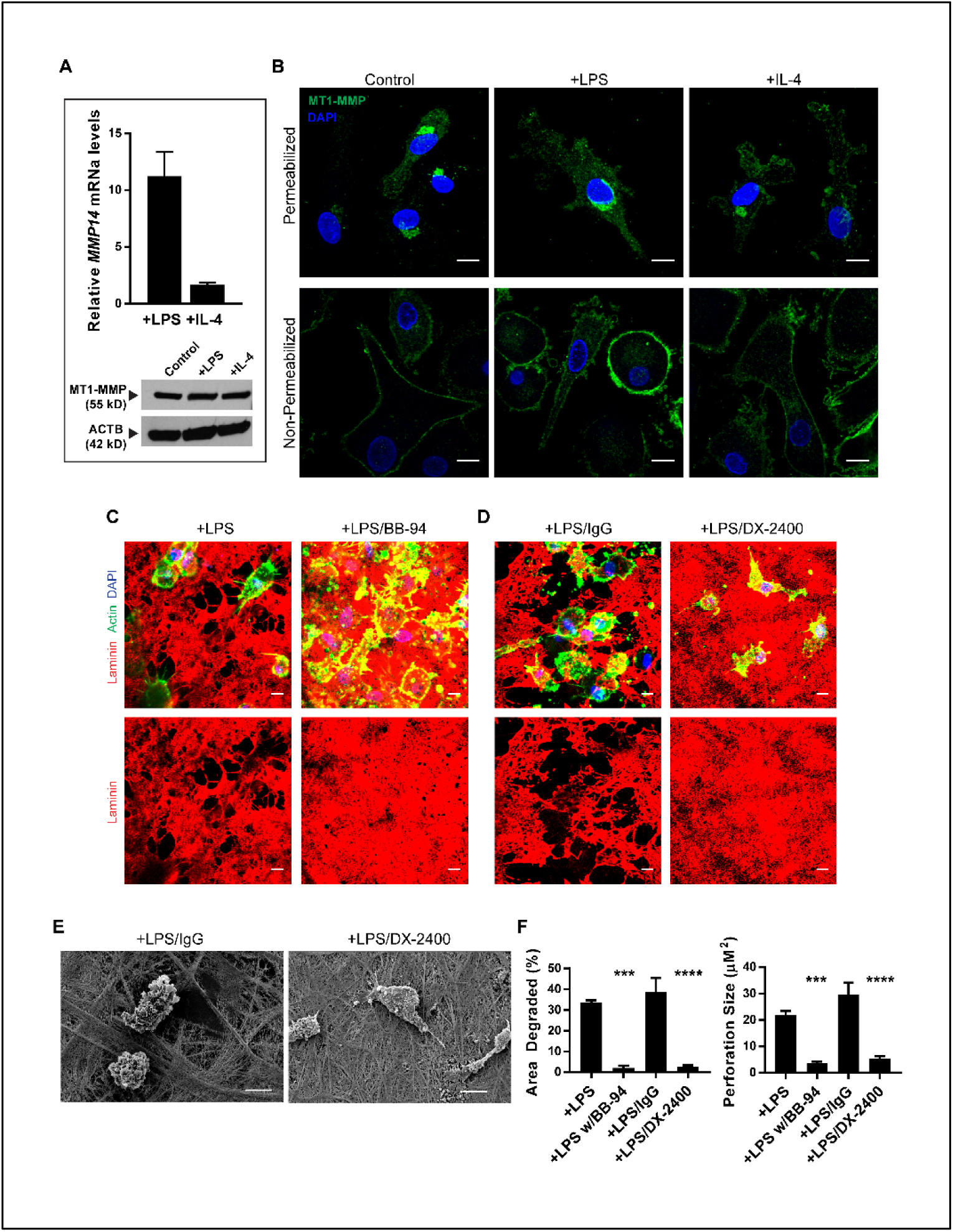
Human macrophages require MT1-MMP to degrade the basement membrane. **(A)** Relative MT1-MMP expression in human macrophages left unstimulated, polarized with LPS (1 μg/mL), or recombinant human IL-4 (20 ng/mL) as determined by qPCR (top panel) or Western blot (bottom panel). **(B)** High-magnification confocal images of endogenous MT1-MMP immunofluorescence in permeabilized (top 3 panels) or non-permeabilized (bottom 3 panels) human macrophages. **(C)** Immunofluorescence of macrophages on basement membrane constructs in the presence of LPS (1 μg/mL) without or with 5 μm BB-94, **(D)** 75 μg/mL IgG control antibody or 75 μg/mL of MT1-MMP blocking antibody, DX-2400. **(E)** Scanning electron micrograph of mesentery basement membrane after culture with macrophages in the presence of LPS (1 μg/mL) and either 75 μg/mL IgG or 75 μg/mL DX-2400 for 6 days. Images shown in (B-D) are representative of three replicates. Bars: (B-E) 10 μm. **(F)** Quantification of the area of basement membrane degraded as analyzed by ImageJ pixel analysis of each condition from (C-D). Results are expressed as mean ± SEM; (****) P < .00001, (***) P < .0001; n = 3.

### Macrophages can traverse the BM–interstitial matrix interface independently of proteolysis

While multiple normal as well as neoplastic cell populations degrade native tissue barriers as a prerequisite for supporting tissue-invasive activity (Hanahan & Weinberg, 2011; Rowe & Weiss, 2009; Wolf et al., 2013), the mechanisms underlying the ability of macrophages to cross native tissue interfaces remains undefined. Confirming the barrier properties of our *ex vivo* model, the highly invasive human breast cancer cell line, MDA-MB-231 (Ota, Li, Hu, & Weiss, 2009; Sabeh et al., 2004; Sabeh, Shimizu-Hirota, et al., 2009), rapidly degrades the underlying basement membrane barrier and infiltrates the interstitial space (Figure 7A,E). By contrast, when MDA-MB-231 cells are cultured in the presence of BB-94, basement membrane remodeling is curtailed and invasion into the interstitium completely blocked (Figure 7A,E). To determine whether macrophage invasion is similarly linked to basement membrane proteolysis, wild-type and MT1-MMP-targeted mouse or human macrophages were cultured atop explants for 6 d and infiltration monitored. As expected, both mouse and human macrophages degrade the subjacent membrane coincident with the expression of tissue-invasive activity (Figure 7B). Interestingly, macrophages accessing the interstitial matrix are found to adhere tightly to elastin fibrils while also infiltrating basement membrane sleeves that ensheathed the peritoneal vasculature (Figure 7C-D). However, in contrast to the protease-dependent invasion program deployed by carcinoma cells, when basement membrane remodeling by macrophages is blocked by targeting MT1-MMP, macrophages continue to infiltrate the interstitial matrix, bind to elastin fibers and invade vascular basement membranes (Figure 7B-E). Similar, if not identical, results are obtained when macrophages are cultured in the presence of protease inhibitors directed against each of the major proteinase classes in tandem, ruling out the possibility that alternate proteolytic systems required for invasion are engaged following MT1-MMP targeting (Figure 7B,E).

**Figure 7.**
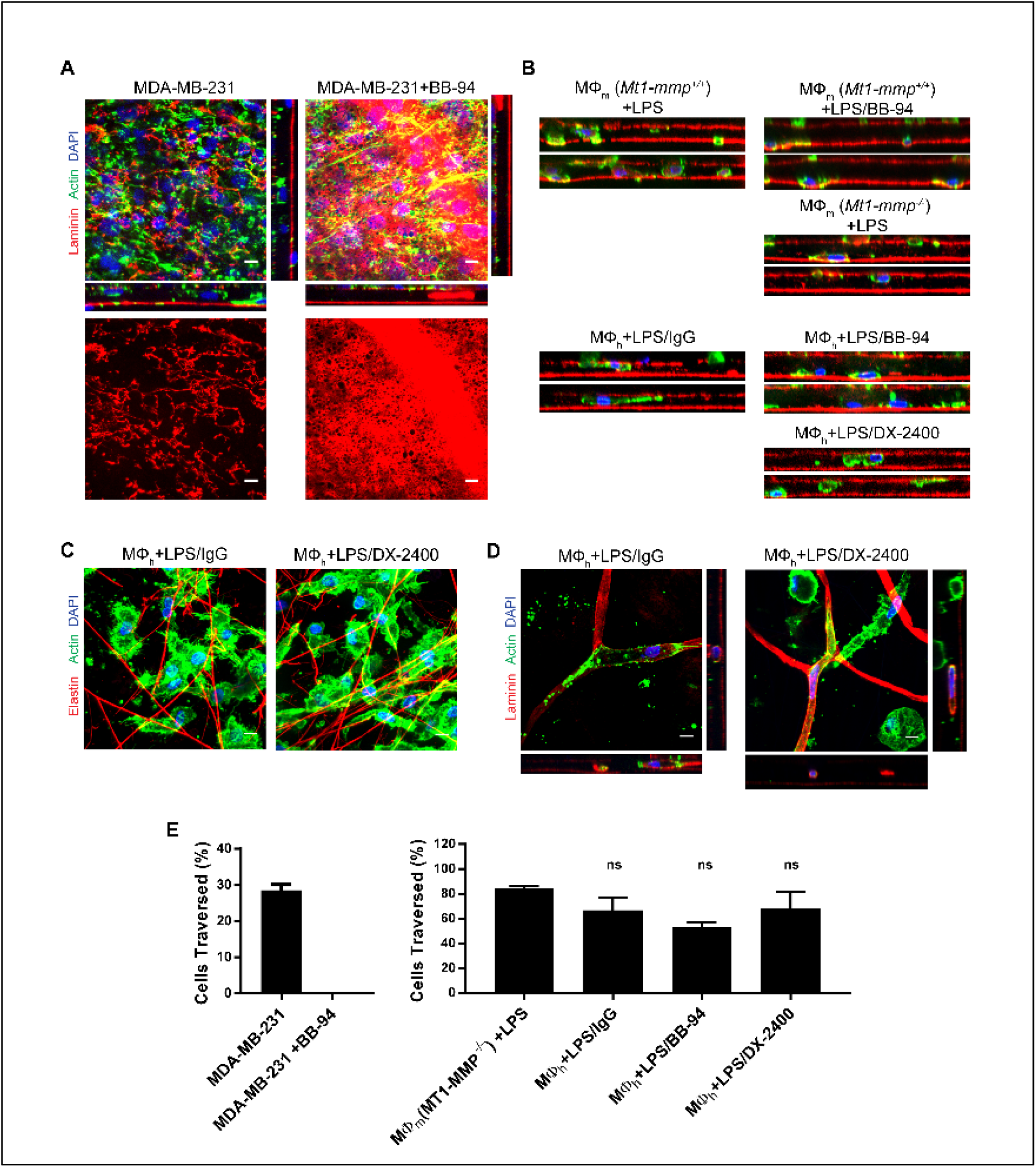
Macrophages can mobilize a proteinase-independent tissue invasion program. **(A)** *En face* and orthogonal immunofluorescence of basement membranes cultured with human MDA-MB-231 breast carcinoma cells for 48 h without or with 5 μm BB-94. **(B)** Orthogonal view reconstructions of LPS (1 μg/mL)-polarized MT1-MMP^+/+^ or MT1-MMP^−/−^ mouse BMDM in the absence or presence of 5 μm BB-94 (top 3 panels), and LPS (1 μg/mL)-polarized human macrophages in the presence or absence of 75 μg/mL IgG, 5 μm BB-94, 75 μg/mL DX-2400, or a protease inhibitor mix (100 μm E-64d, 100 μg/ml aprotinin, 10 μm pepstatin A, 100 μg/ml SBTI, 5 μm BB-94, 2 μm leupeptin) (bottom 4 panels). Images shown are representative of three replicates. **(C)** Immunofluorescence of human macrophages polarized with LPS (1 μg/mL) adhering to the interstitial elastin network in the presence or absence of 75 μg/mL IgG or 75 μg/mL DX-2400 for 6 days. **(D)** *En face* and orthogonal immunofluorescence of human macrophages infiltrating laminin-stained vascular basement membranes in the presence or absence of 75 μg/mL IgG or 75 μg/mL DX-2400 for 6 days. **(E)** Quantification of cell bodies (including nuclei) of MDA-MB-231 (left panel) or macrophages (right panel), located between the two basement membrane layers from (A-B) as a percentage of the total number of cells. Bars: (A, C-D) 10 μm.

### Basement membrane pores provide macrophages with proteinase-independent access to the interstitium

While carcinoma cells mobilize proteinases to degrade the pericellular ECM in order to invade the interstitial matrix, the question remains as to the means by which macrophages traverse an identical barrier independently of proteolytic remodeling. Interestingly, earlier reports have described basement membrane “pores” that exist in almost every tissue where they have been proposed allow for epithelial-mesenchymal or mesodermal-stromal contact – and possibly, myeloid cell trafficking (Barreiro et al., 2016; Bluemink, van Maurik, & Lawson, 1976; Howat, Holmes, Holgate, & Lackie, 2001; Oakford et al., 2011; Takahashi-Iwanaga, Iwanaga, & Isayama, 1999; Takeuchi & Gonda, 2004). As such, we considered the possibility that macrophages might gain access to the interstitium through similar structures via a non-proteolytic process (Howat et al., 2001; McClugage, Low, & Zimny, 1986; Oakford et al., 2011; Pflicke & Sixt, 2009; Toner, Carr, Ferguson, & Mackay, 1970). Indeed, under higher resolution, the peritoneal basement membrane can be shown to harbor a series of ~1 μm diameter pores (Figure 8A). Importantly, these pores are visible in intact tissue prior to decellularization, ruling out pore formation as an unintended consequence of tissue processing (Supplemental Figure 1). Hence, basement membranes and primary human macrophages were fluorescently pre-labeled and transmigration captured by live imaging in the presence of the MT1-MMP-blocking antibody (Figure 8B). Over a 7-hour time-course, macrophages were observed moving in a 2-dimenstional orientation towards a group of preformed portals (Figure 8B, Video 7). Orthogonal reconstructions over this timeframe demonstrate that macrophages first move laterally towards these perforations in an MT1-MMP-independent fashion before migrating vertically through the basement membrane and into the interstitial matrix (Figure 8B, Video 8). Hence, macrophages - unlike carcinoma cells - do not require MT1-MMP to penetrate the basement membrane-interstitial interface or invade the underlying interstitial matrix.

**Figure 8.**
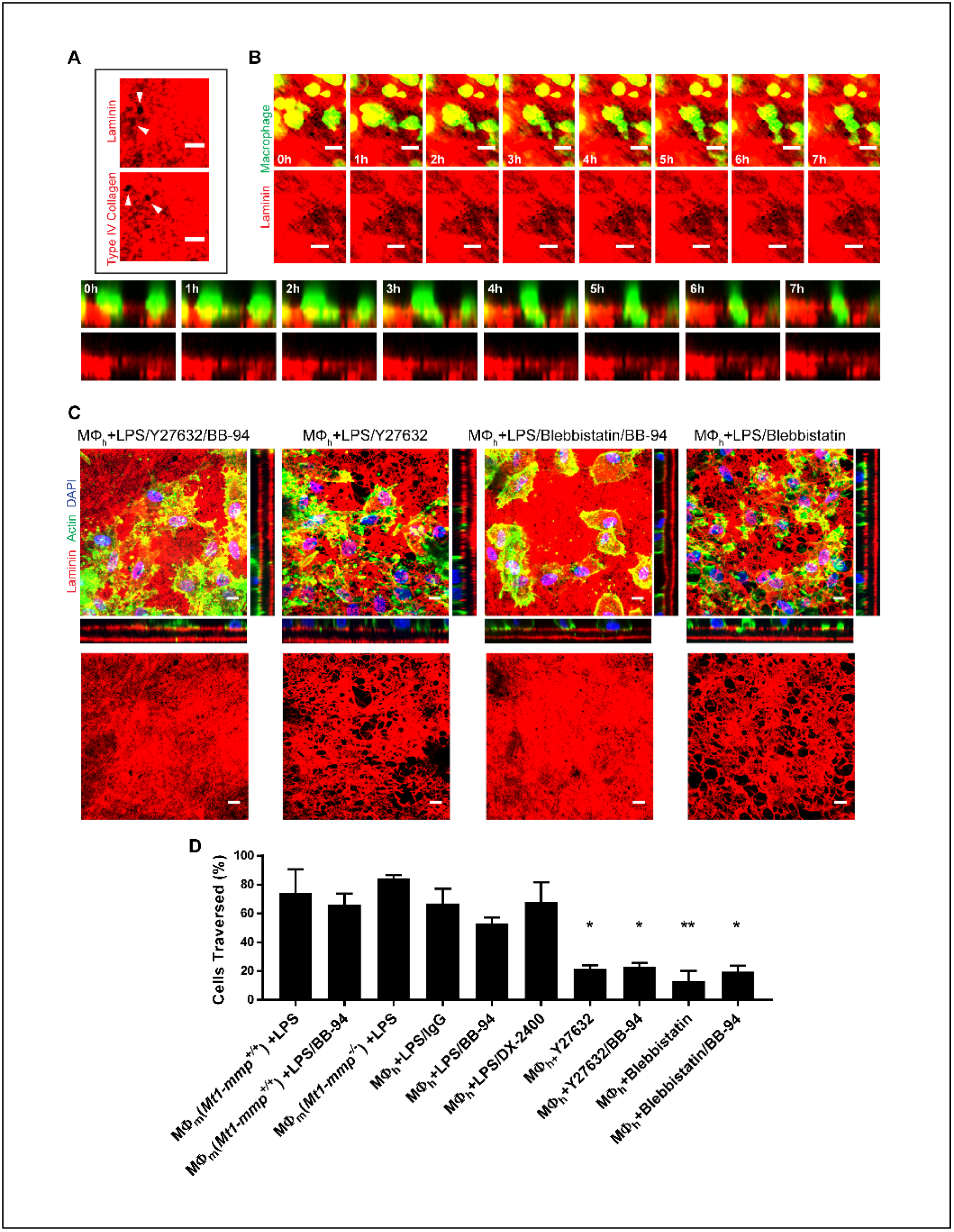
Macrophages traverse preformed portals in the basement membrane in an actomyosin-dependent fashion. **(A)** Magnified confocal immunofluorescence of pores (arrowheads) in laminin- and type IV collagen-labeled basement membranes (B) Time-lapse series of CFSE-labeled human macrophages (green) and laminin-prelabeled basement membrane (red) captured every hour for 7 h immediately after plating. A macrophage moves towards preformed portals and distorts its shape (top two rows) while traversing the basement membrane en route to the interstitial matrix (bottom two rows). **(B)** *En face* and orthogonal immunofluorescence of basement membranes cultured with human macrophages polarized with LPS (1 μg/mL) in the absence or presence of Y-27632 (20 μM) and BB-94 (5 μm) (left panels), or Blebbistatin (20 μM) and BB-94 (5 μm) (right panels). Images shown are representative of three replicates. **(C)** Quantification of cell bodies (including nuclei) of macrophages, located between the two basement membrane layers (from Figure 7B and Figure 8B) as a percentage of the total number of cells. Bars: (A, B) 5 μm, (C) 10 μm. Results are expressed as mean ± SEM; (**) P < .001, (*) P < .01; n = 3.

In other cell systems, non-proteolytic mechanisms of invasion have been linked to the transfer of mechanical forces from the cell body to either the surrounding matrix or the perinuclear compartment as a means to shape the rigid nucleus to a size that allows small ECM pores to be negotiated (Chang et al., 2017; Harunaga, Doyle, & Yamada, 2014; Hung et al., 2013; Paul, Mistriotis, & Konstantopoulos, 2017; Ruprecht et al., 2015). In an effort to define the contribution of actomyosin-dependent contractility to macrophage invasion, human macrophage infiltration into the interstitial matrix was assessed in the presence of the Rho kinase inhibitor, Y27632, or the myosin II inhibitor, blebbistatin (Harunaga et al., 2014; Hung et al., 2013; Ruprecht et al., 2015). While human macrophages cultured in the presence of BB-94 actively cross the basement membrane and infiltrate the interstitial matrix, the addition of either Y27632 or blebbistatin significantly blocks invasion (Figure 8C,D), highlighting the importance actomyosin-dependent forces in supporting motility responses though physiologic tissue barriers. To next determine if proteolysis might generate larger pore sizes in the ECM that potentially preclude a requirement for the actomyosin network, macrophages were cultured atop the ex vivo construct with either Y27632 or blebbistatin; but in the absence of BB-94 (Figure 8C,D). Interestingly, neither inhibitor affects the ability of macrophages to proteolytically remodel the underlying basement membrane (Figure 8C,D). Nevertheless, interstitial matrix invasion remains inhibited, demonstrating a continued requirement for actomyosin-dependent forces as the macrophages traverse the exposed interstitial ECM compartment, presumably as a consequence of negotiating restrictive pore sizes in this matrix compartment as well. Hence, these data are consistent with a heretofore undescribed dual ability of macrophages to infiltrate interstitial matrix by either proteolytically degrading the basement membrane barrier or alternatively, mobilizing actomyosin-dependent pathways to non-proteolytically traverse basement membrane portals.

## Discussion

In postnatal states, macrophage patrol or infiltrate host tissue where they remodel the ECM in order to either exert palliative effects, e.g., during host defense and wound repair, or participate in deleterious outcomes, e.g., chronic inflammatory disease states and cancer (Nathan & Ding, 2010; Noy & Pollard, 2014; Wynn & Vannella, 2016). However, the precise mechanisms that allow macrophages to remodel native tissue barriers have remained largely undefined. To date, almost all studies have relied on the use of model systems to characterize macrophage interactions with either basement membrane or interstitial matrix barriers (Cougoule et al., 2012; Fleetwood et al., 2014; Gui et al., 2014; Guiet et al., 2011; Guiet et al., 2012; Jevnikar et al., 2012; M. Y. Murray et al., 2013; Ogura, Bridgeman, & Malanchi, 2017; Starnes et al., 2014; Van Goethem et al., 2010; Werb, Bainton, & Jones, 1980). However, given increased appreciation that these constructs cannot recapitulate the more complex structure of the ECM *in vivo*, and that the composition and mechanical properties of the ECM can each affect cell function (Liu et al., 2015; Previtera & Sengupta, 2015; Wiesner et al., 2014), the utility of these systems for predicting macrophage function *in vivo* is subject to debate. For example, whereas basement membranes *in vivo* are type IV collagen-rich and mechanically rigid as a consequence of lysyl oxidase- and peroxidasin-mediated covalent crosslinks, *in vitro* constructs that rely on EHS carcinoma extracts (i.e., Matrigel) are alternatively enriched with laminin, mechanically soft and largely devoid of the critical type IV collagen crosslinks that define basement membrane structure (Halfter et al., 2015; Rowe & Weiss, 2008, 2009; Willis, Sabeh, Li, & Weiss, 2013). Likewise, given the fact that the interstitial matrix, though dominated by type I/III collagen, is comprised of hundreds of distinct components, attempts to recapitulate its structure with relatively simple collagen hydrogels is problematic (Naba et al., 2016; Sabeh, Shimizu-Hirota, et al., 2009; Wolf et al., 2013). While these systems as well as synthetic polyethylene glycol- or alginate- based substrates engineered in 2-D, 3-D or microchannel format can nevertheless yield valuable insights (Hung et al., 2013; Liu et al., 2015; Paul et al., 2017; Previtera & Sengupta, 2015; Thiam et al., 2016), none of these constructs recapitulate the structural complexity or architecture of native tissues. Given these limitations, we selected a matrix explant model for characterizing cell-matrix interactions, thereby allowing us to gauge the ability of non-polarized as well as polarized macrophages to remodel and transmigrate native basement membrane barriers while gaining access to an underlying interstitial matrix.

In considering the potential proteolytic mechanisms that might be responsible for macrophage-dependent basement membrane remodeling, we chose an unbiased transcriptional screen as a means to identify candidate proteinases. Consistent with reports implicating cysteine proteinases, serine proteinases as well as secreted MMPs in conferring macrophages with the ability to invade Matrigel-based constructs, each of these proteolytic systems were expressed by polarized macrophages (Jevnikar et al., 2012; M. Y. Murray et al., 2013). However, when macrophage interactions with native basement membranes were examined, targeting these proteinases with class-specific inhibitors failed to block matrix remodeling. Instead, both human and mouse macrophages deployed the membrane-anchored MMP, MT1-MMP, as the dominant effector of basement membrane remodeling. Interestingly, in a manner similar to that observed during podosome-mediated proteolysis (Wiesner et al., 2013; Wiesner et al., 2014), MT1-MMP actively trafficked to invasive membrane protrusions as the macrophages penetrated the basement membrane. While we have not examined this exocytotic process in detail, MT1-MMP trafficking most likely involves the RabGTPase-microtubule system previously detailed by Linder and colleagues (Wiesner et al., 2013; Wiesner, Faix, Himmel, Bentzien, & Linder, 2010).

Following MT1-MMP deletion or inhibition, we anticipated that macrophages would be unable to cross the basement membrane or infiltrate the interstitial matrix. While cells can potentially use cytoskeletal-generated forces to displace non-covalently cross-linked ECM fibers (Gjorevski, S. Piotrowski, Varner, & Nelson, 2015; Pflicke & Sixt, 2009; Sabeh, Shimizu-Hirota, et al., 2009), the basement membrane used here is heavily cross-linked by sulfilimine bonds that would be predicted to render the type IV collagen backbone resistant to mechanical displacement (McCall et al., 2014). Further, while migrating cells can negotiate fixed pores whose size exceeds 10% of the nuclear cross-sectional area, the type IV collagen network has been estimated to limit interfibrillar pore size to ~50 nm in diameter, dimensions that would effectively preclude cellular transmigration (Fidler et al., 2017; Hallmann et al., 2015; Kelley et al., 2014; Wolf et al., 2013). Indeed, while human breast carcinoma cells were able to degrade and penetrate the basement membrane, invasion was, as expected (Hanahan & Weinberg, 2011; Hotary et al., 2006; Ota et al., 2009; Paul et al., 2017; Rowe & Weiss, 2008), inhibited completely when MMP activity was blocked. Though recent reports have emphasized the ability of human carcinoma cells, including MDA-MB-231 cells, to alternatively adopt an amoeboid phenotype to negotiate ECM barriers via non-proteolytic mechanisms, these studies consistently rely on artificial matrix models whose relevance to native tissue barriers remains to be determined (Aung et al., 2014; Haeger, Wolf, Zegers, & Friedl, 2015; He & Wirtz, 2014; Liu et al., 2015). Using tissue explants, carcinoma cells were, by contrast, wholly dependent on MMP-dependent proteolysis. Nevertheless, in the absence of MT1-MMP activity, both human and mouse macrophages retained the ability to cross the basement membrane-interstitial matrix interface via a process that is dependent on actomyosin-generated forces. In our efforts to visualize the sites permissive for proteinase-independent transmigration, our attention focused on discrete ~3 μm^2^ pores that decorate the basement membrane surface (Figure 8A, arrowheads; Supplemental Figure 1). Importantly, micrometer-sized basement pores have been identified in lung, skin, blood vessel and colon tissues, raising the distinct possibility that these structures are purposefully generated during embryogenesis not only to allow epithelial/mesodermal-stromal crosstalk, but also to serve as permissive passageways for cell movement (Howat et al., 2001; McClugage et al., 1986; Oakford et al., 2011; Pflicke & Sixt, 2009; Toner et al., 1970). Given their relatively small pore size - at least relative the nuclear dimensions of most cell populations - the engagement of the macrophage actomyosin network is consistent with recent studies demonstrating similar requirements as cells negotiate space-restrictive environments (Hung et al., 2013; Paul et al., 2017; Thiam et al., 2016). Interestingly, although the basement membrane surrounding lymphatic vessels are less well-organized than those found subtending epithelial cells or the vascular endothelium, Sixt and colleagues recently demonstrated that basement membrane portals similar to those described here are permissive for non-proteolytic dendritic cell trafficking (Pflicke & Sixt, 2009). It should be stressed however, that basement membrane pores do not provide proteinase-independent access to all cell populations, as exemplified by the inability of MDA-MB-231 cells to usurp these passageways in their efforts to access the interstitial compartment. Furthermore, the trafficking mechanisms outlined here for macrophages cannot be extended to all myeloid population as we have found that human neutrophils are unable to cross the peritoneal basement membrane – either in the absence or presence of chemotactic gradients (unpublished observation). While Glentis et al have recently proposed that cancer-associated fibroblasts can induce cancer cell invasion through the peritoneal basement membrane by a metalloproteinase-independent process (Glentis et al., 2017), SEM images of their constructs appear to be devoid of an intact basement membrane and only show the fibrillar matrix of interstitial collagen (i.e., the diameter of type IV collagen fibrils is only on the order of 2 nm and requires high resolution metal shadow casting for visualization) (Yurchenco & Ruben, 1987). As the ability of cell-generated mechanical forces to guide carcinoma cells through stromal collagen hydrogels has been described (Aung et al., 2014), their model is most consistent with the ability of cancer-associated fibroblasts to accelerate cancer cell invasion through a porous collagen network. These issues notwithstanding, macrophages may well represent a unique cell population that can traverse basement membranes in a cell autonomous fashion by alternatively mobilizing MT1-MMP-dependent or proteinase-independent mechanisms. In this regard, we note that recent live imaging studies of macrophage migration within zebrafish larvae describe a dual requirement for proteinases and ROCK-dependent traction forces (Barros-Becker, Lam, Fisher, & Huttenlocher, 2017). However, as a cocktail of proteinase inhibitors was used in this study, the identity of the targeted proteinases remains to be determined.

Having gained access to the interstitial matrix, we noted that macrophages were not randomly arrayed within the stroma, but instead were closely associated with the underlying network of elastin fibrils. Interestingly, inflammatory macrophage infiltrating adipose tissues *in vivo* were recently reported to be similarly positioned, raising the possibility that elastin networks may provide a 1-dimensional ‘highway’ for macrophage trafficking through stromal tissues (Martinez-Santibanez et al., 2015). Of note, we also observed that macrophage-associated elastin fibrils were frequently fragmented (Figure 7C). As elastin networks were disrupted in the absence or presence of MT1-MMP, alternate proteolytic systems must be in play here. Indeed, we have previously described the ability of human macrophages to degrade insoluble elastin fibers by mobilizing the cysteine proteinases, cathepsin L and S, and the role of these proteinases in this more physiologic model of elastinolytic activity remains to be determined (Filippov et al., 2003; Punturieri et al., 2000; Reddy et al., 1995). Finally, we also note that macrophages, having gained access to the interstitial matrix, establish contact with the stromal face of both vascular basement membranes and the inner face of the reflected basement membrane. Recent studies demonstrate that basement membranes are not homogenous, but instead are asymmetrically organized with marked differences in rigidity and matrix composition on the epithelial versus stromal sides, raising the possibility that macrophage-basement membrane trafficking might be unidirectionally favored (Halfter, Monnier, et al., 2013). However, we find that macrophages are able to transmigrate basement membranes in either direction, a finding consistent with recent reports demonstrating the ability of macrophages to remodel vascular basement membranes during carcinoma cell intravasation or the alveolar basement membrane following lung injury (Harney et al., 2015; Misharin et al., 2017; Wyckoff et al., 2007).

In sum, we find that human as well as mouse macrophages mobilize MT1-MMP as the dominant effector of basement membrane remodeling. While MT1-MMP confers non-polarized as well as polarized macrophages with the ability to resorb native tissues during tissue trafficking, these cells can also adopt an alternate phenotype that allows them to traffic through tissue barriers in a proteinase-independent mode that is not conserved in carcinoma cells. As such, we posit that macrophages, unlike other normal or neoplastic cell populations, have the ability to infiltrate tissues wherein the ECM is purposefully left unscathed during reparative states or irreversibly remodeled in association with tissue-destructive events.

## Materials and Methods

### Isolation of primary macrophages

Bone marrow macrophages were isolated as previously described from 2–8 week old wild-type *(Mt1-mmp^+/+^)* or MT1-MMP-null *(Mt1-mmp^−/−^)* Swiss Black mice (Holmbeck et al 1999; Sakamoto and Seiki 2009). Briefly, long bones were flushed with PBS, red cells were lysed with ACK buffer (ThermoFisher) and the remaining cells were cultured in alpha-MEM with 10% heat-inactivated fetal bovine serum (HI-FBS), 1% penicillin-streptomycin solution (ThermoFisher), and 10 ng/mL M-CSF (R&D Systems) overnight on tissue culture dishes. Nonadherent cells were plated onto non-tissue culture-treated dishes in media with M-CSF for an additional 5–7 days; media was replaced every 48 hours.

Human peripheral blood monocytes were isolated from whole blood of volunteers in accordance with institutional review board (IRB) approval and the patient’s informed consent. PBMCs were separated by Lymphocyte Separation Medium (Corning) by density centrifugation, purified by CD14 selection (Miltenyi Biotec) and cultured at 2 × 10^6^ in 6-well plates containing RPMI 1640 without serum. After 2 h, media was replaced with RPMI 1640 with 1% penicillin-streptomycin solution and 20% autologous serum for 5–7 days. Autologous serum was prepared by incubating non-heparinized whole blood at 37 °C for 1 h followed by centrifugation at 2,850 g for 15 minutes, and sterile filtration of the serum fraction.

### *Ex vivo* mesentery ECM preparation

Mesentery tissue were prepared as previously described (Witz et al., 2001; Hotary et al., 2006). Briefly, rat mesentery was mounted on 6.5 or 12-mm diameter Transwells (Sigma) with sterile surgical thread and decellularized with 0.1 N ammonium hydroxide. 1-2 × 10^5^ mouse or human macrophages were cultured atop the tissue for six days with media changes every 48 hours. All experiments were performed in complete medium in the absence or presence of the following inhibitors 100 μM E-64d, 100 μg/mL aprotinin, 100 μg/mL soybean trypsin inhibitor (SBTI), 20 μM Y-27632, 20 μM Blebbistatin (Sigma), 5 μM BB-94 (Tocris Bioscience), 12.5 μg/mL TIMP-1, 5 μg/mL TIMP-2 (Peprotech). Protease inhibitor mix contained 100 μM E-64d, 100 μg/mL aprotinin, 100 μg/mL SBTI, 5 μM BB-94, 2 μM leupeptin, 10 μM pepstatin A. Human macrophages were also cultured with 75 μg/mL human isotype control IgG antibody or anti-MT1-MMP antibody DX-2400 (Ager et al., 2015) in medium with 20% heat-inactivated autologous human serum and 1% penicillin-streptomycin in the presence of 5 μL Fc-receptor blocking antibody TruStain FcX (Biolegend). DX-2400 was provided by the Kadmon Corporation. Macrophages were polarized with 1 μg/mL LPS from *Escherichia coli* O111:B4 (Sigma) or 20 ng/mL recombinant mouse or human IL-4 (Peprotech). After six days of culture, tissue constructs were washed with PBS, fixed with 4% PFA, and stained as described.

### Lentiviral gene transfer

A mCherry-tagged MT1-MMP construct (Sakurai-Yageta 2008) was cloned into pLenti lox IRES EGFP vector and subsequently transfected into 293T cells using Lipofectamine 2000 (ThermoFisher) to generate lentiviral particles. BMDM at 5 days post-isolation were incubated with the lentivirus-containing supernatant in the presence of 8 μg/mL polybrene for 6 h before media was replaced. 48 h later transduced macrophages were cultured atop the tissue construct as described.

### Tumor cell culture

MDA-MB-231 were cultured in DMEM supplemented with 10% heat-inactivated fetal bovine serum (HI-FBS) and a 1% penicillin-streptomycin solution. Cells were cultured on the basement membrane construct for 48 hours before processing.

### Confocal fluorescence microscopy and analysis

PFA-fixed constructs were incubated with polyclonal antibodies targeting laminin (Sigma, cat #: L9393), type IV collagen (Abcam, cat #: ab19808), and elastin (EMD Millipore cat #: 2039), at 1:150 dilution in a blocking solution of 1% bovine serum albumin-PBS for 1 h room temperature. Constructs were then incubated with secondary fluorescent antibodies at 1:250 dilution while cells were labeled with Alexa Fluor 488 phalloidin and DAPI (Sigma) for 1 h in blocking solution. Image acquisition was performed using a spinning disc confocal CSU-WI (Yokogawa) on a Nikon Eclipse TI inverted microscope with a 60x oil-immersion objective and the Micro-Manager software (Open Imaging). Fluorescent images were processed with ImageJ (National Institutes of Health) with 3D viewer plugin for orthogonal and 3D reconstructions. Confocal imaging of the collagen I matrix was captured by second harmonic generation on a Leica SP5 inverted confocal microscope with a 60x oil-immersion objective.

For immunofluorescence of endogenous MT1-MMP, fixed primary human macrophages were incubated on glass coverslips with 1:50 rabbit monoclonal anti-MT1-MMP (Abcam) overnight at 4°C in 3% BSA-PBS with or without 0.1% Triton X-100 to permeabilize the cells, followed by incubation with 1:200 Alexa Fluor-488 secondary antibody for 1 h 37°C.

### Live image microscopy

Live imaging was performed on unfixed tissue constructs pre-labeled with fluorescent antibodies as above. Macrophages were incubated with 5 μM CFSE (Life Technologies) in PBS for 20 minutes at 37°C, quenched with a 5x volume of medium with 1% HI-autologous serum, resuspended in PBS-Fc receptor block for 5 minutes at room temperature, and plated on the pre-labeled tissue construct. Z-stacks or single slices were captured in a 37°C 5% CO_2_ humidified chamber (Livecell Pathology Devices) as described. Cell outlines were generated and overlaid using the binary and outline functions of ImageJ.

### Electron microscopy

Tissue constructs were processed for SEM as follows, fix in 2% glutaraldehyde/1.5% paraformaldehyde in 0.1 M cacodylate buffer, post-fix in 1% osmium tetroxide, and dehydrated through a graded ethanol series as described (Hotary et al. 2003). Image acquisition was performed using an AMRAY 1910 field emission scanning electron microscope at 5.0 kV.

### ELISA

Anti-Rat COL4A1 ELISA kits were purchased from LSBio. Tissue constructs were cultured for 72 hours in the presence of media alone or with LPS-stimulated human macrophages and LPS before cell-free media was analyzed according to the manufacturer’s instructions.

### qPCR and transcriptional profiling

RNA was isolated from macrophages using the NucleoSpin RNA kit (Macherey Nagel) as instructed. Day 5–7 macrophages were polarized for 24 hours as described. cDNA synthesis was performed with Superscript III enzyme (Invitrogen). qPCR reactions were performed in triplicate with SYBR green PCR master mix on a 7900HT fast Real-Time PCR machine (Applied Biosystems). Data were analyzed using the comparative threshold cycle method with mRNA levels normalized to GAPDH.

For transcriptional profiling, total mRNA was isolated as above, and labeled and hybridized to Mouse Gene ST 2.1 strips (Affymetrix). Three replicates of each sample were analyzed by the University of Michigan Microarray Core. BMDM proteases with expression values greater than 2^4^ in any condition were tabulated and further analyzed for relative fold differences across conditions.

### Western Blot

Western blots were performed as described with antibodies targeting MT1-MMP (Epitomics), alpha 2 (IV) NC1, Clone H22 (Chondrex), and β-actin (Cell Signaling). Primary antibodies were labeled with horseradish peroxidase-conjugated species-specific secondary antibodies (Santa Cruz) and detected by the SuperSignal West Pico system (Pierce). For type IV collagen dimer-monomer content analysis, isolated tissue was first digested with bacterial collagenase type IV (Worthington Biochemical Corporation) overnight at 37 °C with occasional vortexing. Samples were pelleted at 15,000 g for 20 minutes and analyzed by SDS-PAGE without heat-denaturation.

### Statistical Analysis

The area of basement membrane degraded as well as basement membrane perforation size were calculated using ImageJ as follows; image intensity was enhanced and background subtracted using default settings, the images converted to black and white via binary function, inverted, and particles larger than a 1 μm^2^ minimum cut off analyzed. For percent invasion, cells were considered traversed if the cell body including nucleus were located between the two basement membrane layers. Results are expressed as mean ± SEM and comparisons were made with one-way ANOVA.

## Acknowledgements

We thank the Kadmon Corporation for providing the anti-MT1-MMP antibody, licensed to the Kadmon Corporation by the Dyax Corporation. We acknowledge Stephanie King (Life Sciences Institute) for assistance with illustrations and Craig Johnson (University of Michigan) for assistance with microarray analysis. We thank Stefan Linder for helpful discussions. This work was supported by grants from the NIH to SJW (AI105068) and the Cancer Biology Training Grant (NCI Training Grant T32-CA009676).

The authors declare no competing financial interests.

## Author contributions

J.C. Bahr and S.J. Weiss conceptualized and designed the experiments, analyzed the data, wrote and edited the manuscript. J.C. Bahr carried out the experiments and formal analysis.

---

Supplemental Figure 1. **Preformed portals in basement membrane prior to decellularization. (A)** *En face* and orthogonal immunofluorescence of mesentery fixed in 4% PFA prior to decellularization. **(B)** Magnification of boxed region in (A) showing a grouping of preformed portals in the basement membrane. Bars: (A, B) 10 μm.

**Video 1**. 3D rotation of rat mesentery construct. Laminin-stained (red) apical and reflected basal basement membrane surfaces in a 360° rotation. Depicted as a 3D-rendered confocal z-stack. Bar 20 μm. Refers to Figure 1B.

**Video 2**. 3D rotation of rat mesentery construct. Elastin (blue) with second harmonic generation of type I collagen (yellow) in a 360° rotation. Depicted as a 3D-rendered confocal z-stack. Bar 20 μm. Refers to Figure 1B.

**Video 3**. Human macrophages actively protruding through the basement membrane. 3D confocal time-lapse view of the interstitium-facing side of the apical basement membrane (red) as human macrophages (green) actively protrude through it. Time of observation: 160 minutes, captured every 10 minutes. Playback: 6 frames/s. Bar 20 μm. Refers to Figure 1H.

**Video 4**. Human macrophages expand basement membrane perforations. Enlarged max intensity projection confocal time-lapse of human macrophages (green) atop the apical basement membrane face (red) in the presence of LPS (1 μg/mL) actively expanding a perforation (arrowhead) in the basement membrane. Time of observation: 160 minutes, captured every 10 minutes. Playback: 6 frames/s. Bar 10 μm. Refers to Figure 1l.

**Video 5**. Human macrophages expand basement membrane perforations. Max intensity projection and orthogonal reconstruction confocal time-lapse of human macrophages (green) atop the apical basement membrane face (red) in the presence of LPS (1 μg/mL) actively expanding a perforation in the basement membrane (arrowhead). Time of observation: 11 h, captured every 1 h. Playback: 6 frames/s. Bar 10 μm. Refers to Figure 2.

**Video 6**. Human macrophages expand basement membrane perforations. Max intensity projection confocal time-lapse combined with static confocal orthogonal reconstructions of human macrophages (green) atop the apical basement membrane face (red) in the presence of LPS (1 μg/mL) actively expanding a perforation in the basement membrane (arrowhead). Time of observation: 90 minutes, captured every 3 minutes. Playback: 10 frames/s. Bar 10 μm. Refers to Figure 2.

**Video 7**. Human macrophages traverse preformed basement membrane portals. Enlarged max intensity projection confocal time-lapse of human macrophages (green) atop the apical basement membrane face (red) in the presence of LPS (1 μg/mL) and 75 μg/mL DX-2400. Time of observation: 460 minutes, captured every 20 minutes. Playback: 6 frames/s. Bar 5μm. Refers to Figure 8A.

**Video 8**. Human macrophages traverse preformed basement membrane portals. Enlarged orthogonal reconstruction confocal time-lapse of human macrophages (green) atop the apical basement membrane face (red) in the presence of LPS (1 μg/mL) and 75 μg/mL DX-2400. Time of observation: 520 minutes, captured every 20 minutes. Playback: 8 frames/s. Refers to Figure 8A.

